# Propofol directly binds and inhibits skeletal muscle ryanodine receptor 1 (RyR1)

**DOI:** 10.1101/2024.01.10.575040

**Authors:** Thomas T. Joseph, Weiming Bu, Omid Haji-Ghassemi, Yu Seby Chen, Kellie Woll, Paul D. Allen, Grace Brannigan, Filip van Petegem, Roderic G. Eckenhoff

## Abstract

As the primary Ca^2+^ release channel in skeletal muscle sarcoplasmic reticulum (SR), mutations in the type 1 ryanodine receptor (RyR1) or its binding partners underlie a constellation of muscle disorders, including malignant hyperthermia (MH). In patients with MH mutations, exposure to triggering drugs such as the halogenated volatile anesthetics biases RyR1 to an open state, resulting in uncontrolled Ca^2+^ release, sarcomere tension and heat production. Restoration of Ca^2+^ into the SR also consumes ATP, generating a further untenable metabolic load.

When anesthetizing patients with known MH mutations, the non-triggering intravenous general anesthetic propofol is commonly substituted for triggering anesthetics. Evidence of direct binding of anesthetic agents to RyR1 or its binding partners is scant, and the atomic-level interactions of propofol with RyR1 are entirely unknown. Here, we show that propofol decreases RyR1 opening in heavy SR vesicles and planar lipid bilayers, and that it inhibits activator-induced Ca^2+^ release from SR in human skeletal muscle. In addition to confirming direct binding, photoaffinity labeling using *m-*azipropofol (AziP*m*) revealed several putative propofol binding sites on RyR1. Prediction of binding affinity by molecular dynamics simulation suggests that propofol binds at least one of these sites at clinical concentrations. These findings invite the hypothesis that in addition to propofol not triggering MH, it may also be protective against MH by inhibiting induced Ca^2+^ flux through RyR1.

## Introduction

Ryanodine receptor 1 (RyR1) is the primary Ca^2+^ release channel in the sarcoplasmic reticulum (SR) of skeletal muscle. It is a critical element of excitation-contraction coupling, together with the voltage-gated calcium channel Ca_V_1.1, STAC3, and Junctophilin-1 or Junctophilin-2 (1). A complex interplay of allosteric mechanisms controls the opening of RyR1, including small molecules, protein binding partners, and post-translational modifications (2).

Dysregulation of RyR1 is implicated in the pathophysiology of a constellation of muscle disorders. These include central core disease (3,4), multiminicore disease (5), and malignant hyperthermia (MH) (6). During an MH episode, RyR1 is biased to the open state, resulting in uncontrolled flow of Ca^2+^ ions out of the SR, causing heat production and often muscle rigor. Returning this large excess of Ca^2+^ to the SR also consumes ATP. This hypermetabolic state results clinically in acidosis, hyperkalemia, rhabdomyolysis, and hyperthermia. The necessary conditions for MH are 1) a causative mutation in RyR1, Ca_V_1.1, STAC3, or a yet-to-be identified additional gene and 2) the presence of a triggering drug, such as volatile anesthetics or succinylcholine. Even when all these factors are present, an MH episode is not guaranteed. It is presumed, but not directly shown, that triggering drugs bind to RyR1 and/or its partners, biasing the channel to an open state.

Hundreds of disease-associated mutations in RyR1 have been reported, of which a subset has been enumerated as causative by the European Malignant Hyperthermia Group (EMHG, https://www.emhg.org). Many MH mutations are found at domain-domain interfaces (7,8), and cryo-EM structures of a disease causing mutant RyR1 have shown that the mutations result in different conformations linked to channel opening, by destabilizing the closed, resting state of the channel (9–11).

When MH is diagnosed in an anesthetized patient, anesthesia must often be continued to permit a safe conclusion to the surgical procedure. The administration of triggering agents is discontinued, and a non-triggering anesthetic begun. Propofol, a GABA_A_ receptor agonist, is the general anesthetic of choice for maintence under this circumstance because it has not been reported to trigger MH (12,13). In transfected HEK-293 cells with MH-mutant RyR1, propofol did not promote caffeine-induced Ca^2+^ release (14). Overall, propofol has not previously been shown to affect RyR1-mediated calcium release at clinically tractable concentrations (15,16).

In this manuscript, we describe a combination of approaches providing evidence that propofol binds to and inhibits RyR1 opening at clinically tractable concentrations. First, we show that propofol and its photoaffinity analogue *m*-azi-propofol (AziP*m*) decrease [^3^H]ryanodine binding to SR vesicles and RyR1 channel open probability, both in wild-type and R615C mutant RyR1. We confirm the expected consequence on ryanodine-induced Ca^2+^ release from cultured skeletal muscle. Next, we identify several propofol binding sites through analogue photoaffinity labeling (17–19). Finally, we computationally predict the binding affinity of propofol to one of the identified binding pockets using streamlined alchemical free energy perturbation molecular dynamics (SAFEP MD) (20,21), suggesting that propofol may be bound to this site at clinically tractable concentrations. Taken together, our results suggest that propofol binds and inhibits RyR1 opening at clinical concentrations and may thereby prevent the clinical manifestations of MH even with exposure to triggering agents such as the volatile anesthetics.

## Results

### AziP*m* and propofol each decrease the proportion of open wild-type RyR1s as measured by ryanodine binding

We used a tritiated ryanodine assay, which reflects the proportion of open RyR1s, to measure RyR1 opening as a function of the concentration of AziP*m* and propofol in heavy sarcoplasmic reticulum (SR) vesicles (22). We tested 0–48 µM AziP*m* and 0–100 µM propofol with RyR1. At low Ca^2+^ concentrations (pCa 7, 100 nM), AziP*m* and propofol each reduced the proportion of open channels, with IC_50_ values of 6.9 µM and 5.8 µM respectively. At activating Ca^2+^ concentrations (pCa 5, 10 µM), a similar trend was observed, with IC_50_ values of 4.8 µM and 6.7 µM respectively (Figure 1).

**Figure 1.**
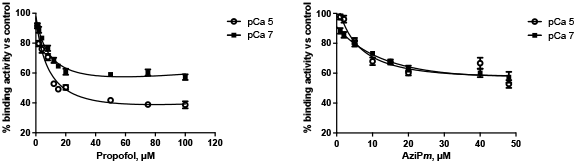
[^3^H]ryanodine binding to skeletal muscle heavy SR vesicles as a function of propofol (left) or AziPm concentration (right). Values expressed as percentage of control (0 µM propofol or AziPm)

### AziP*m* and propofol inhibit R615C RyR1 channel opening in planar lipid bilayers

We measured the channel open probability of homozygous pig R615C RyR1 reconstituted in planar lipid bilayers (PLBs) as a function of the concentrations of AziP*m* and propofol, in the presence of an activating concentration of Ca^2+^ (40 µM). The R615C mutation predisposes pigs to the MH-like porcine stress syndrome (23), and therefore can serve as a model for propofol’s influence on RyR1 with an MH mutation; the analogous human MH-causative RyR1 mutation is R614C. In the control experiment with neither AziPm nor propofol, channel open probability was 0.11. This decreased, respectively, to 0.03, 0.07, and 0.04 with 10 µM AziPm, 10 µM propofol, and 30 µM propofol. (Figure 2).

**Figure 2.**
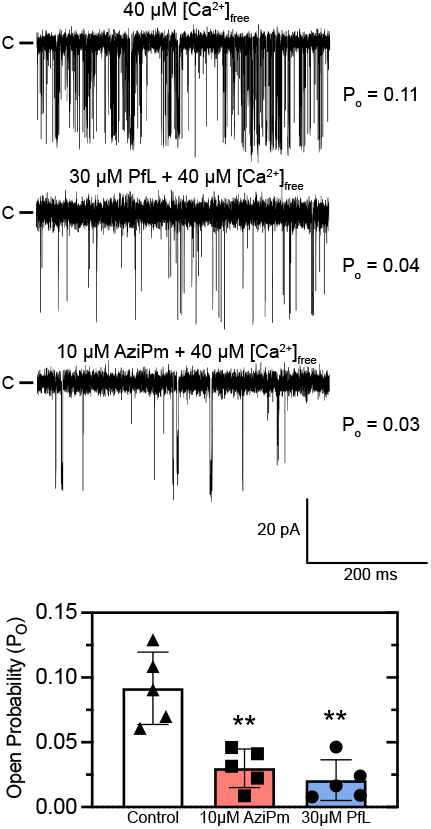
RyR1 channel opening probability as a function of propofol and AziP*m* concentration in PLBs. ** statistically differenent from control (n = 5, p < 0.01, Mann-Whitney test)

### Propofol inhibits activator-mediated RyR1 channel opening in wild-type cultured human skeletal myotubes (HSM)

Using Fura-2 Ca^2+^ imaging, we studied the effect of propofol on intracellular Ca^2+^ concentration in wild-type HSM in the presence of the RyR1 activator ryanodine. Higher measured 340:380 nm ratios indicate increased Ca^2+^ release, attributable to RyR1 opening. Without propofol, the 340:380 nm ratio saturated at a ryanodine concentration of approximately 1–3 µM (Figure 3). This enabled us to select maximally potent concentrations of activators to test the effect of propofol.

**Figure 3.**
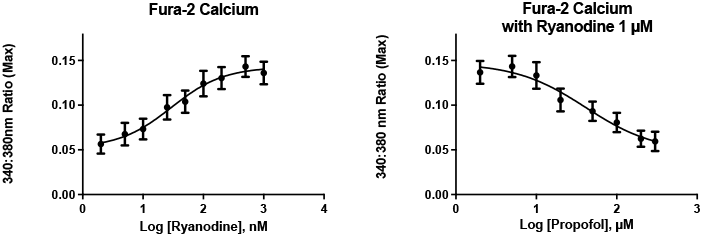
340:380 nm ratio with activators and ryanodine, as a function of propofol concentration. Greater open probability with higher ryanodine concentration is implied (left). At a constant concentration of ryanodine, increasing propofol concentration results in lower open probability.

When propofol was added to HSM/Fura-2 preparations containing 1 µM ryanodine, the 340:380 nm ratio decreased as propofol concentration increased, suggesting that propofol inhibits RyR1 opening in the presence of this activator. The mean IC_50_ of propofol in the presence of 1 µM ryanodine was 29.5 µM (SE: 0.1 µM).

### AziP*m* binding sites identified on RyR1 by photoaffinity labeling

As the photoaffinity ligand AziP*m* is chemically similar and has similar anesthetic effects to propofol (17), it is likely that residues photoadducted by AziP*m* would correspond to propofol binding sites. Photoaffinity labeling with AziP*m* was conducted with three variants of RyR1 purified, respectively, from skeletal muscle of wild-type rabbit (*Oryctolagus cuniculus*), wild-type pig (*Sus scrofa*), and RyR1-R615C pig. Rabbit and pig RyR1 have 97% sequence identity with human RyR1, with most non-identical residues residing outside the transmembrane domain.

For each experiment, a preparation of RyR1 and FKBP12.6 (calstabin-2, which stabilizes the RyR1 closed state) was incubated with AziP*m* (17). After irradiation resulting in the formation of covalent AziP*m* adducts, the photoadducted RyR1 residues were identified using mass spectrometry. The sequence coverage was 83%, 87.5%, and 80.2% across the rabbit, pig WT, and pig RyR1-R615C proteins respectively.

To evaluate whether AziP*m* binds to sites that propofol does not (*i*.*e*., false positives), a competition (or protection) assay was conducted. Pretreatment with 200 µM propofol prevented adduction by AziP*m*, implying that the set of AziP*m* binding sites is a subset of propofol binding sites. Taken together with the planar-lipid bilayer results showing that both ligands decrease RyR1 channel open probability, these data suggest that AziP*m* and propofol bind substantially the same locations on RyR1.

The identified sites are listed in Table 1 and depicted in Figure 4. Below, we refer to all photoadducted sites by their sequence locations in the rabbit RyR1 unless present only in pig RyR1, or as otherwise noted. Many of the sites are adjacent to RyR1 regions of functional significance. Several S2-S3 linker mutations have previously been shown *in vitro* to confer RyR2-like Ca^2+^-dependent regulation on RyR1, where the channel open probability as measured by [^3^H]ryanodine binding is increased at higher, activating Ca^2+^ concentration (24). The caffeine binding site was previously visualized by cryo-EM as a pocket including E4239, W4716, and I4996 (25).

**Table 1.**
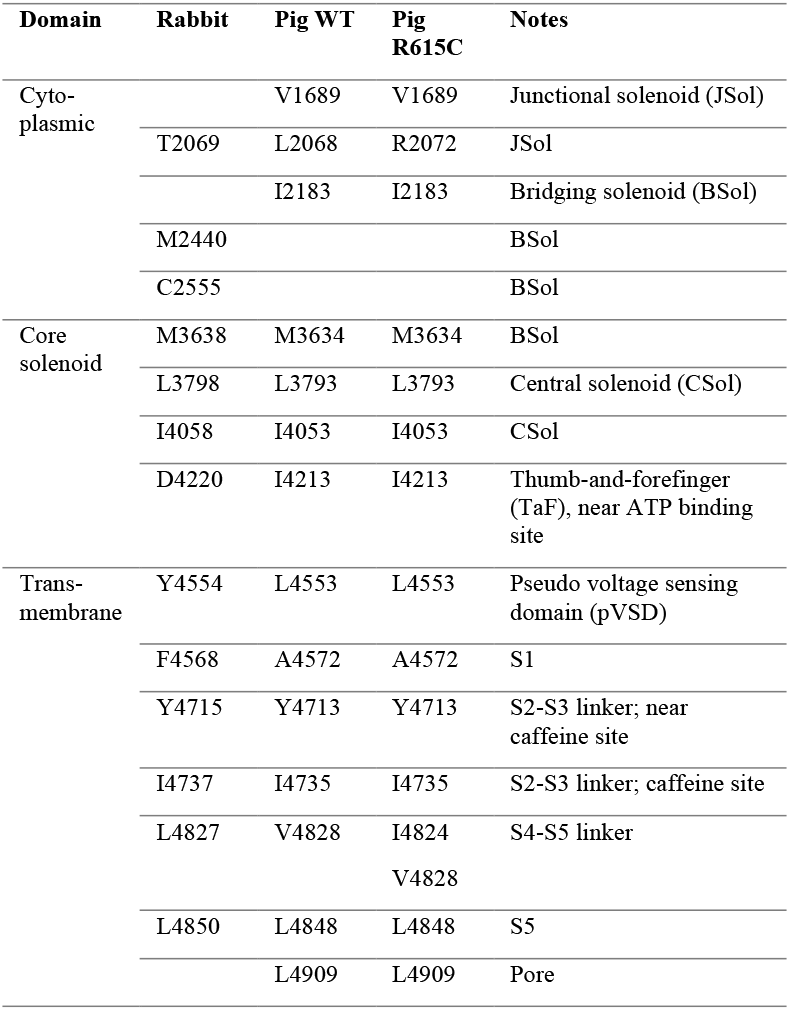
AziP*m* photoadducted sites in RyR1.

**Figure 4.**
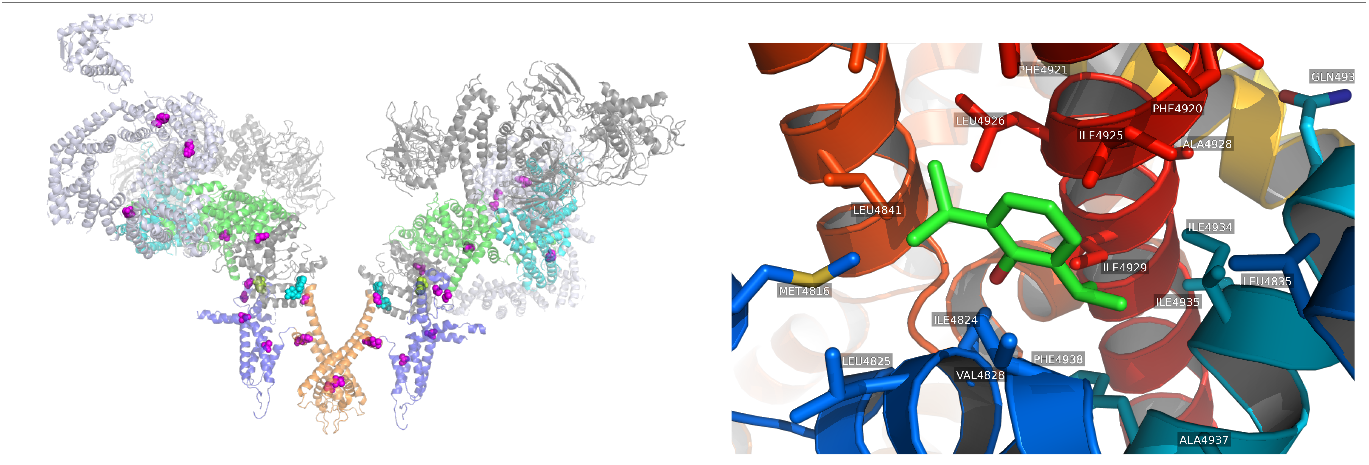
Left pane: RyR1 residues photoadducted by AziP*m*, in magenta spheres. For visual clarity only two of the four monomers are shown. For reference, bound ATP is shown in cyan spheres, caffeine in red spheres. JSol in cyan, Bsol in light blue, Csol in green, pVSD in blue, pore in orange. Right pane: V4828 site (right, closer to pore) occupied by propofol. Representative pose taken from equilibrium MD simulations.

The L4827 residue in the S4-S5 linker forms part of a binding pocket surrounded by lipophilic α-helices. This intersubunit binding pocket is part of a region adjacent to the pore lumen (Fig. 1). Mutations in this region (T4825I, H4832Y in rabbit RyR1) increase Ca^2+^ affinities for activation and decrease it for deactivation *in vitro*: channel open probabilities are increased at both low and high Ca^2+^ concentrations, but not at intermediate concentrations (24).

### Propofol binds proximally to V4828 with predicted affinity similar to clinical concentration

Since only the photoadducted residue is identified instead of the specific adducted atom (*e*.*g*. side chain or carbonyl) the bound pose of the non-photolyzed parent ligand is not known. To predict the pose of the parent ligand in one of the adducted pockets, we conducted all-atom equilibrium MD simulation on the transmembrane domain of RyR1 with the parent ligand placed in the V4828 pocket, followed by FEP MD simulations to predict the binding affinity.

We chose the pocket adjacent to V4828 in pig RyR1 (V4830 in rabbit RyR1). This was prioritized for a computationally intensive analysis because of its proximity to the RyR1 pore (~5 Å in the plane perpendicular to the pore axis, measured between the Cα atoms of L4827 and pore-lining residue I4937), and would have a shorter physical pathway to modify pore opening probability.

Because it would be computationally prohibitive to sample an informatively large subset of the configurational space of the entirety of RyR1, we studied only the pore-containing transmembrane domain (TMD, residues 4546–5029) embedded in a lipid bilayer surrounded by water with 0.15 M KCl. As docking algorithms are not well adapted to hydrophobic, low-affinity molecules, we initialized the system by manually placing a ligand molecule in the pocket. Minimization and equilibration MD simulation included a flat-bottomed volume restraint on the ligand center of gravity in order to prevent escape of the low-affinity ligand from the site. This approach, previously used in equilibrium MD simulations of isoflurane with a two-pore potassium channel (26), imposes no energetic penalty when the center of geometry of the ligand remains within a 5 Å radius sphere in the pocket. This was used at this stage to prevent escape of the ligand from the region before locating an energetic minimum. We then ran 50 ns of production MD. The open-state R615C pig RyR1 was used, although the R615C mutation was not included in the simulation since it is outside of the transmembrane domain.

In production MD simulation, the ligand remained in a stable orientation. The propofol hydroxyl group remained oriented away from V4828 and toward the pore. We used double-decoupling streamlined alchemical free energy perturbation (21) (SAFEP) to predict the binding affinity of the ligand, with the ligand in the aqueous phase serving as the unbound reference point. SAFEP calculations predicted the Gibbs free energy of propofol binding in the V4828 site 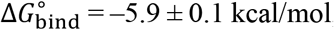, corresponding to *K*_D_ = 55.8 µM (95% CI: 40.3–77.3 µM). This includes aqueous phase decoupling energy 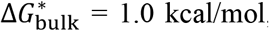, restraint correction ∆*G*_DBC_ = 0.9 ± 0.1 kcal/mol, and volumetric correction ∆*G*°= 0.4 kcal/mol. For the FEP calculations, convergence was excellent (details in Supp. Fig. S1). Of note, since there is a logarithmic dependence of *K*_D_ on ∆G, small changes in ∆G in this range can lead to large changes in *K*_D_.

## Discussion

In this study, we 1) showed that propofol decreases RyR1 open probability in planar lipid bilayers and human skeletal muscle cells, 2) showed that propofol inhibits [^3^H]-ryanodine binding to SR vesicles, 3) identified by photoaffinity labeling several propofol binding sites on RyR1, and 4) calculated binding affinity of propofol in two selected photoadducted sites adjacent to each other and near the pore in WT RyR1. Overall, our results show that propofol both binds RyR1 in specific sites and inhibits pore opening.

The plasma concentration of propofol at loss of consciousness in human volunteers was estimated to be on the order of 10 µM (27). We found that the IC_50_ for RyR1 inhibition in HSM was within an order of magnitude of this. Since propofol avidly binds serum proteins (28), the plasma concentration required to elicit RyR1 inhibition would be higher as compared to that in an otherwise protein-free environment as with our HSM experiments. Moreover, as propofol is highly lipid soluble, its concentration can be expected to be amplified in lipid-rich environments such as protein transmembrane domains. Propofol given as the primary anesthetic may therefore actively mitigate the clinical manifestations of MH, or other disease states arising from increased RyR1 activation. Though we calculated propofol binding affinities only for two binding pockets, we found that corresponding dissociation constants were in the µM range, suggesting that these are relevant at clinical concentrations.

In each of its four subunits, the RyR1 TMD has at least 6 transmembrane helices of which four encode a (pseudo-)voltage-sensing domain, similar to the inositol-3,4,5-triphosphate (IP3) receptor and voltage-gated ion channels in the Na_v_, K_v_, and Ca_v_ families. Propofol inhibition of channels with TMDs similar to that of the RyR1 is not a unique phenomenon (29–34). Moreover, binding in or around the conserved S4-5 linker domain appears to be a canonical feature in ion channels such as NaChBac, Na_v_Ms, and K_v_1.2 (35– 38). Given the existence of multiple propofol binding sites distributed among different domains in this very large protein, we hypothesize that an allosteric mechanism at least partly underlies its effects, although this is conjectural. One possibility is disruption of subunit cooperativity, shown in a kinetic model to be important to the function of the structurally similar RyR2 (39).

There are some limitations to our study. It is difficult to assess the sensitivity of photoadduction for binding sites of the parent ligand: *e*.*g*. AziP*m* might adduct sites that the parent ligand propofol does not bind, or propofol may bind sites for which we did not detect AziP*m* adduction. This is both because the diazirine group and halogens render it chemically distinct from the parent ligand, and because less than 100% of the RyR1 sequence was covered. However, at least in apoferritin, photolabeling, crystallography and fluorescence competition placed AziP*m* and propofol in the same site (17). As we did not evaluate a functional model of MH using intact muscle, our prediction that propofol inhibits the clinical presentation of MH is a hypothesis to be tested in the future. Nonetheless, from a photochemistry perspective it is gratifying to note that identical or highly analogous residues were adducted in RyR1 purified from three different sources, and that no apparent photochemical selectivity for specific residues was observed.

Because of the availability of pig R615C RyR1 (40), we also compared photoadduction to wild-type and R615C pig RyR1. Despite substantial global conformational changes induced by the mutation (9) the photoadducted residues were the same with one exception (L2068 in WT *versus* R2072 in R615C). This suggests that the presence of the R615C mutation increases the likelihood of RyR1 activation, rather than altering anesthetic binding, although again, not all binding sites may have been detected.

Our results may be transferable to other disease processes involving RyRs. For example, calcium dysregulation thought to be a central aspect of the pathogenesis of Alzheimer disease. As all three subtypes, RyR1, RyR2, and RyR3, of the ryanodine receptors are present in neuronal endoplasmic reticulum, RyR channel dysfunction is correlated to disease progression (41,42). This invites the hypothesis that a propofol-based anesthetic is less likely to aggravate Alzheimer disease, than, for example, known triggering anesthetics like isoflurane. Some clinical evidence is consistent with this hypothesis (41–43). Hence, a propofol-based general anesthetic is a rational choice both from the perspective of neurodegeneration as well as MH risk.

## Materials and Methods

### [^3^H]Ryanodine binding assay

The [^3^H]ryanodine binding assay was carried out as previously described (22). Heavy SR (HSR) vesicles were provided by Dr. Francisco Alvarado (Cardiovascular Research Center, School of Medicine and Public Health, University of Wisconsin, Madison, WI). The binding assays were performed at fixed [Ca^2+^] = 100 nM (pCa 7) or 10 μM (pCa 5).

Briefly, propofol (1–100μM) or AziP*m* (1–48 μM) was added to a binding mixture with 100 μl volume, containing 50μg of HSR protein, 200 mM KCl, 100 mM HEPES buffer (pH 7.2), 5 nM [^3^H]ryanodine (56 Ci/mmol, PerkinElmer, #NET950), 1 mM EGTA and enough CaCl_2_ to set free [Ca^2+^] at 100 nM (pCa 7) or 10 µM (pCa 5). The Ca^2+^/EGTA ratio for these solutions was calculated using MaxChelator (WEBMAXC Lite V1.15; temperature=37 °C, pH 7.2, Ionic=0.3N). Non-specific binding was determined under the same conditions in the presence of 20 μM unlabeled ryanodine (Millipore-Sigma, #559276).

Binding reactions were incubated for 2 hours at 37°C. Prior to the reaction, the filters were presoaked with 0.5% polyethylenimine for 30 mins. Then the reaction was stopped by vacuum filtration and washed three times with the distilled water. The filters were soaked in 10 mL Ecolite(+)™ scintillation cocktail overnight. [^3^H]Ryanodine binding was measured by liquid scintillation in a PerkinElmer Tri-Carb 2800TR counter. Experiments were done in triplicate.

### Purification of rabbit or pig RyR1 from rabbit/pig muscle

Approximately 200 g of frozen rabbit/pig skeletal muscle was blended for 120 s in 2 × 500 ml of 10 mM tris/maleate (pH 6.8), 10% sucrose, 1 mM dithiothreitol (DTT), 1 mM EDTA, 200 μM PMSF, and 1 mM benzamidine. The mixture was centrifuged at 4°C for 10 min at 7000g. The supernatant was filtered through a cheesecloth and centrifuged at 4°C for 40 min at 40,000g. Pellets were solubilized in buffer S1 containing 20 mM Hepes (pH 7.4), 1 M KCl, 2 mM TCEP, 150 μM PMSF, 1 mM EGTA, 1% CHAPS, and 0.2% soybean phosphatidylcholine with 100 μl of protease inhibitor cocktail (Protease Inhibitor Cocktail Set III, EDTA-Free, Calbiochem). After stirring for 1 hour at 4°C, the solubilized membranes were diluted with 120 ml of buffer S2 (as for buffer S1 but lacking 1 M KCl). His-GST-FKBP12.6 (~5 mg) was then added to the solubilized membranes, with incubation for 2 hours at 4°C. After ultracentrifugation for 45 min at 200,000g, the supernatant was filtered and mixed with 2 to 3 ml of GS4B resin (Cytiva) pre-equilibrated with buffer S3 containing 20 mM HEPES (pH 7.5), 0.5 M KCl, 0.5% CHAPS, 0.2% soybean phosphatidylcholine, 1 mM EGTA, 2 mM TCEP, 150 μM PMSF, and 100 μl of protease inhibitor cocktail, with stirring for 3 hours at 4°C. The mixture was poured into a column and washed with 10-CV buffer S3. Next, 10 ml of buffer S4 (identical to S3 except with 75 mM HEPES) together with 1.5 mg of TEV protease was added to the column and incubated overnight to elute RyR1 from the column. The eluents were concentrated to 500 μl, applied to a gel-filtration column (Superose 6 10/300 GL, Cytiva) and eluted with buffer S5 containing 15 mM Tris (pH 7.5), 0.15 M NaCl, 0.5% CHAPS, 0.001% 18:1 (Δ9-cis) phosphatidylcholine (DOPC), 2 mM TCEP, and 1 mM EGTA. Peak fractions containing RyR1 complexes were combined and concentrated to ~ 2 mg/ml. Concentration was estimated using NanoDrop [absorbance at 280 nm, 1% (w/v) = 1.0].

### RyR1 proteolipsome reconstitution

RyR1 was reconstituted into proteoliposomes as previously described (9). Briefly, a 5:3 mixture of 1,2-dioleoyl-sn-glycero-3-phosphoethanolamine (DOPE) and 1,2-dioleoyl-sn-glycero-3-phosphocholine (DOPC) (Avanti Polar Lipids) were dried into a thin film, followed by overnight incubation in a vacuum chamber. Dried lipids (1 mg) were solubilized with 400 μl of rabbit RyR1 (0.7 mg/ml) in buffer S5, and the mixture was dialyzed overnight using a 3.5-kDa membrane in 10 mM HEPES (pH 7.4), 500 mM NaCl, 0.1 mM EGTA, 0.2 mM CaCl_2_, 0.15 mM PMSF, and 1 mM DTT. Following dialysis, the samples were aliquoted, flash-frozen in liquid nitrogen, and stored at −80°C for later use.

### Planar lipid bilayer methods: single channel recordings

For all experiments, planar lipid bilayers were formed from a mixture of DOPE:DOPC= 5:3 (mol/mol) dissolved in decane with a final lipid concentration of 20 mg/mL. For single channel recordings, an integrated chip-based recording setup Orbit mini and EDR2 software (Nanion Technologies, Livingston, NJ) was employed. Recordings were obtained in parallel with multielectrode-cavity-array chips (Ionera Technologies, Freiburg, Germany). The cis and trans chambers contained symmetrical solutions of 250 mM HEPES, 150 mM KCl, 1 mM EGTA (pH 7.3), 0.2 mM CaCl_2_ ([Ca^2+^] free = 0.1 μM). The free concentration of Ca^2+^ was calculated with WebMaxC program (v.2.50; https://somapp.ucdmc.ucdavis.edu/pharmacology/bers/maxchelator/webmaxc/webmaxcS.htm). Ca^2+^ concentrations were further verified using PerfectIon™ Combination Calcium Electrode unit (Mettler Toledo). To promote fusion to prepared suspended bilayers, 5% glycerol was incorporated into the proteoliposomes. In brief, prepared proteoliposomes were mixed 1:1 in 250 mM HEPES (pH 7.3), 150mM KCl, and 10% glycerol. The samples were then freeze-thawed and sonicated 3-5 times to incorporate the glycerol into the proteoliposome lumen. 1–2 μL RyR1 proteoliposomes were added to the cis chamber. To further promote fusion the voltage was maintained at +40 mV. Recordings were started at the point of successful RyR1 insertion. Propofol was introduced by the displacement of 3–30% of the cis chamber volume with recording solution containing 60 μM or 100 μM propofol. Concentrations of propofol aqueous stocks were determined by measuring the absorbance at 270 nm, using ε 1584 M^-1^ cm^-1^ (44). All RyR1 measurements were conducted at 22°C and at a constant voltage of –60mV. Recordings were filtered at final bandwidth of 10,000Hz. Clampfit software (10.6, Molecular Devices, San Jose, CA) was used to analyze current traces and only channels with a conductance greater then 700 pS were included in the analysis (45).

### Calcium imaging in human skeletal myotubes

Human skeletal muscle cells (HSMC, Sigma, #150-05F) were maintained in growth medium (Sigma, #151-500) in a 5% CO_2_ atmosphere at 37°C. The medium was changed every other day. Cells were passaged when they reached 80-90% confluence.

To induce differentiation, the cells were plated at 20,000 per cm^2^ on 35 mm dishes (World Precision Instruments, Inc, #FD 35-100) pre-coated with collagen I (Sigma-Aldrich, Louis, MO, USA) at 37 °C and 5% CO2. The cells were incubated overnight in growth medium, then the growth medium was replaced with differentiation medium (Sigma, #151D-250). The differentiation medium was changed every other day. Multinuclear myotubes typically formed within 5-6 days.

### Calcium fluorescence measurements

Differentiated myotubes were washed with the running HBSS buffer (Hepes-buffered salt solution (HBSS, Sigma, #H-4891) containing 1.8 mM CaCl_2_, 0.8 mM MgCl_2_).

The cells were loaded with 1.5 μM Fura-2 AM (Invitrogen, #F1221) and 20% BSA in the running HBSS buffer for 45 min. The cell was washed again and incubated with 150 µL running buffer for 15 min to allow de-esterification of the acetoxymethyl ester from the now-intracellular Fura-2 AM. The cells were then ready to be excited alternately at 340 nm and 380 nm. The fluorescent maker Fura-2 binds to intracellular Ca^2+^, with the ratio of emission due to excitation at 340 nm and 380 nm directly related to the concentration of Ca^2+^. Hence release of calcium stores in the SR into the cytoplasm due to RyR1 channel opening would be expected to result in an increased 340:380 nm ratio (46–48). The ratio of 340nm:380nm was measured using a fluorescent microscope (Olympus IX-70, Japan) equipped with a cooled high-speed digital video camera (Hamamatsu, Japan), and using the software MetaFluor for Olympus (version 7.10.4.407, MetaMorph 2020, Molecular Devices, LLC). 40-50 cells were usually chosen to measure the changes in Fura-2 fluorescence.

#### Experiment 1: Reactivity to ryanodine

Running HBSS buffer containing increasing concentrations of ryanodine (2, 5, 10, 25, 50, 100, 200, 500, 1000 nM; in Figure 3 plotted on a logarithmic scale) were added to the myotubes. Changes in Fura-2 340/380nM ratios were measured.

#### Experiment 2: Reactivity to propofol with 1 µM ryanodine

From the dose response curve for ryanodine we found that 1 µM ryanodine resulted in the maximum Fura-2 340:380 nm ratio, and as a result this concentration was used to determine the inhibition, if any, caused by different concentrations of propofol. The various propofol concentrations (2, 5, 10, 20, 50, 100, 200, 300 µM) was added to 1µM ryanodine. The 340/380 nm ratios were measured using the same method in Experiment 1.

#### Data analysis

To trace the concentration-response curves, the data were normalized to the maximal response of cells. Then, the IC_50_ was calculated from the concentration response curves and all data were analyzed using PRISM 5.0 software (GraphPad Software, San Diego, CA, USA).

### Photolabeling of RyR1-FKBP12.6

A final concentration of 5μM AziP*m* was added (with or without 200 μM propofol) to the purified RyR1-FKBP12.6 to a final protein concentration of 1 μg/μl. The 200 μM propofol was added to test for labeling protection (competition) in order to evaluate AziP*m* binding specificity. The samples were equilibrated on ice in the dark for 5 min and then irradiated for 30 min at 350 nm with an RPR-3000 Rayonet lamp in 1-mm path length quartz cuvettes through a 295-nm glass filter (Newport Corporation).

### In-solution protein digestion

After UV exposure proteins were precipitated overnight at -20° C in 4 volumes of chilled acetone. Protein was pelleted for 20 min at 16,000 x g at 4 °C then gently washed twice with 300 μL of chilled acetone. Protein pellets were air-dried before resuspension in 50 μl of 50 mM Tris-HCl, pH 8.0, 1% Triton X-100, and 0.5% SDS. Insoluble debris was pelleted by centrifugation at 16000x g. The samples were resuspension in final concentration of 50 mM NH_4_HCO_3_. Following 1 μL 0.5 M dithiothreitol (DTT) was added and samples were incubated at 56 °C for 30 min 0.55 M iodoacetamide (IAA) was then added and protein samples were incubated at room temperature in the dark for 45 min. Sequencing grade-modified trypsin (Promega) was added to a final 1:20 protease: protein ratio (w:w) with additional of 0.2% (w/v%) ProteaseMax™ surfactant. Proteins were digested overnight at 37° C. Trypsin digested peptides were diluted to 200 μL with final concentration of 100 mM NH_4_HCO_3_ and 0.02% ProteaseMAX Surfactant prior to the addition of sequencing grade chymotrypsin (Promega) to a final 1:20 protease:protein ratio (w:w). Proteins were digested overnight at 37° C. Acetic acid (AcOH) was added to until the pH < 2 and the peptide digests were incubated at room temperature for 10 min prior to centrifugation at 16000x g for 20 min to remove insoluble debris. The sample was desalted using C18 stage tips prepared in house. Samples were dried by speedvac and resuspended in 0.1% formic acid immediately prior to mass spectrometry analysis.

### In-gel protein digestion

Photolabeled proteins were separated by SDS-PAGE. The identified band corresponding to rRyR1 was excised. Excised bands were distained, dehydrated and dried by speedvac before proteins were reduced by incubation at 56° C for 30 min in 5 mM DTT and 50 mM NH_4_HCO_3_. The DTT solution was removed and proteins were then alkylated by the addition of 55 mM IAA in 50 mM NH_4_HCO_3_ and incubation at room temperature for 45 min in the dark. Bands were dehydrated and dried by speed vac before resuspension in 100 μL 0.2% ProteaseMAX™ surfactant (Promega) and 50mM NH_4_HCO_3_ solution containing trypsin (sequencing grade, Promega) at a 1:20 protease:protein ratio (w:w). Proteins were digested for 12–16 hrs at 37°C. After trypsin digestion, the samples were diluted to 200 μL with final concentration of 100 mM NH_4_HCO_3_ and 0.02% ProteaseMAX™ Surfactant. The samples were further digested overnight at 37 °C with the addition of sequencing grade chymotrypsin (Promega) to a final protease:protein (w:w) ratio of 1:20. Multiple peptide extractions were performed in order to increase hydrophobic peptide retrieval from the gel. The peptide extractions were pooled from the different ratio of acetylnitrile and acetic acid. The final extractions were sonicated for 20 min and dried by speed vac before resuspension in 0.5% acetic acid and further acidified until the pH < 2. Samples were sonicated for 10 min prior to centrifugation at 16 000 x g for 20 min to remove insoluble debris. Samples were desalted using C18 stage tips prepared in house. Samples were dried by speed-vac and resuspended in 0.1% formic acid immediately prior to mass spectrometry analysis.

### Mass spectrometry

Mass spectrometry was performed similarly to the previously reported procedure (49). Briefly, desalted peptides were injected into a Thermo LTQ Orbitrap XL Mass Spectrometer (Thermo Fisher Scientific, Waltham, MA, USA) or an Orbitrap Elite™ Hybrid Ion Trap mass spectrometer. Peptides were eluted with 100 min with linear gradients of ACN in 0.1% formic acid in water (v/v%) starting from 2% to 40% (85 min), then 40% to 85% (5 min) and finally 85% (10 min).

Spectral analysis was conducted using MaxQuant (50) to search b and y ions against the sequence containing rRyR1. All analyses included dynamic oxidation of methionine (+15.9949 m/z) as well as static alkylation of cysteine (+57.0215 m/z; iodoacetamide alkylation). Filter parameters were Xcorr scores (+1 ion) 1.5, (+2 ion) 2.0, (+3 ion) 2.5, deltaCn 0.08, and peptide probability 0.05. Photolabeled peptides were searched with the additional dynamic AziP*m* modifications. Both the in-solution and in-gel sequential trypsin/chymotrypsin digests were searched without enzyme specification with a false discovery rate of 0.01. Samples were conducted in triplicate and samples containing no photoaffinity ligand were treated similarly to control for false positive detection of photoaffinity ligand modifications. To confirm the photolabeled adduct, mass spectrometry work was repeated at the Proteomics Core Facility of the Wistar Institute (Philadelphia, PA, USA).

### Molecular dynamics simulations

We used a structural model determined by cryo-EM, purified from porcine muscle (9). Only the central pore domain of RyR1 – which itself is a functional channel (51) – was simulated, for the sake of computational feasibility. A system containing the entire protein would be prohibitively large. Simulating the whole protein to a timescale where significant global conformational changes could be observed, especially as part of an allosteric mechanism that would influence pore opening from distant binding sites, was computationally impractical, particularly for demanding free energy perturbation MD simulations.

The molecular mechanics parameters for propofol were taken from a previous simulation study of propofol (52). All simulations were conducted with NAMD 2.14 (53), with GPUs used for equilibrium MD simulations. The CHARMM36 force field (54,55) and TIP3P water model were used. Systems were constructed using CHARMM-GUI (56). Production simulations were conducted in the isothermic-isobaric ensemble with Langevin thermostat. The lipid bilayers consisted of 70% 1-palmitoyl-2-oleoyl-sn-glycero-3-phosphocholine (POPC) and 30% cholesterol.

We calculated the absolute binding free energy of propofol 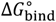. Free energy perturbation (FEP) molecular dynamics simulations were conducted according to the Streamlined Alchemical Free Energy Perturbation (SAFEP) methodology, described in detail in (21). We followed the set-up and analysis procedure described in (20). Briefly, SAFEP uses a limited set of restraints on the ligand to maintain its bound conformation during alchemical transformations and improve sampling of states that most contribute to the binding free energy. The restraints are then corrected for to yield an accurate absolute binding free energy. The overall expression is

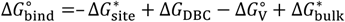

where 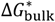 is the energy of decoupling the unbound ligand from solvated to gas phase, 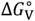 and ∆*G*_DBC,_energies of volumetric and distance-from-bound-conformation (DBC) restraints respectively, and –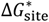 is the energy of coupling the ligand from gas phase to the protein-bound state. 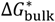 and 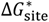 were calculated using FEP MD, ∆*G*_DBC,_ using thermodynamic integration, and 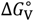 parametrically.

For the calculation of 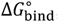, 160 windows were used; for 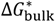, 481 windows; and for ∆*G*_DBC,_, 40 windows. Interleaved double-wide sampling was used. Restraints were imposed using the Colvars module (57). The Bennett Acceptance Ratio method was used to calculate free energy differences in FEP calculations.

## Acknowledgements

We are grateful to Francisco J. Alvarado, University of Wisconsin-Madison, for assistance with the [^3^H]ryanodine binding assay. RGE, FVP and TTJ were supported by National Institute of General Medical Sciences, National Institutes of Health (RGE, FVP, TTJ: R01GM135633, TTJ: K08GM139031).

## Supplementary Information

**Supplementary Figure S1.**
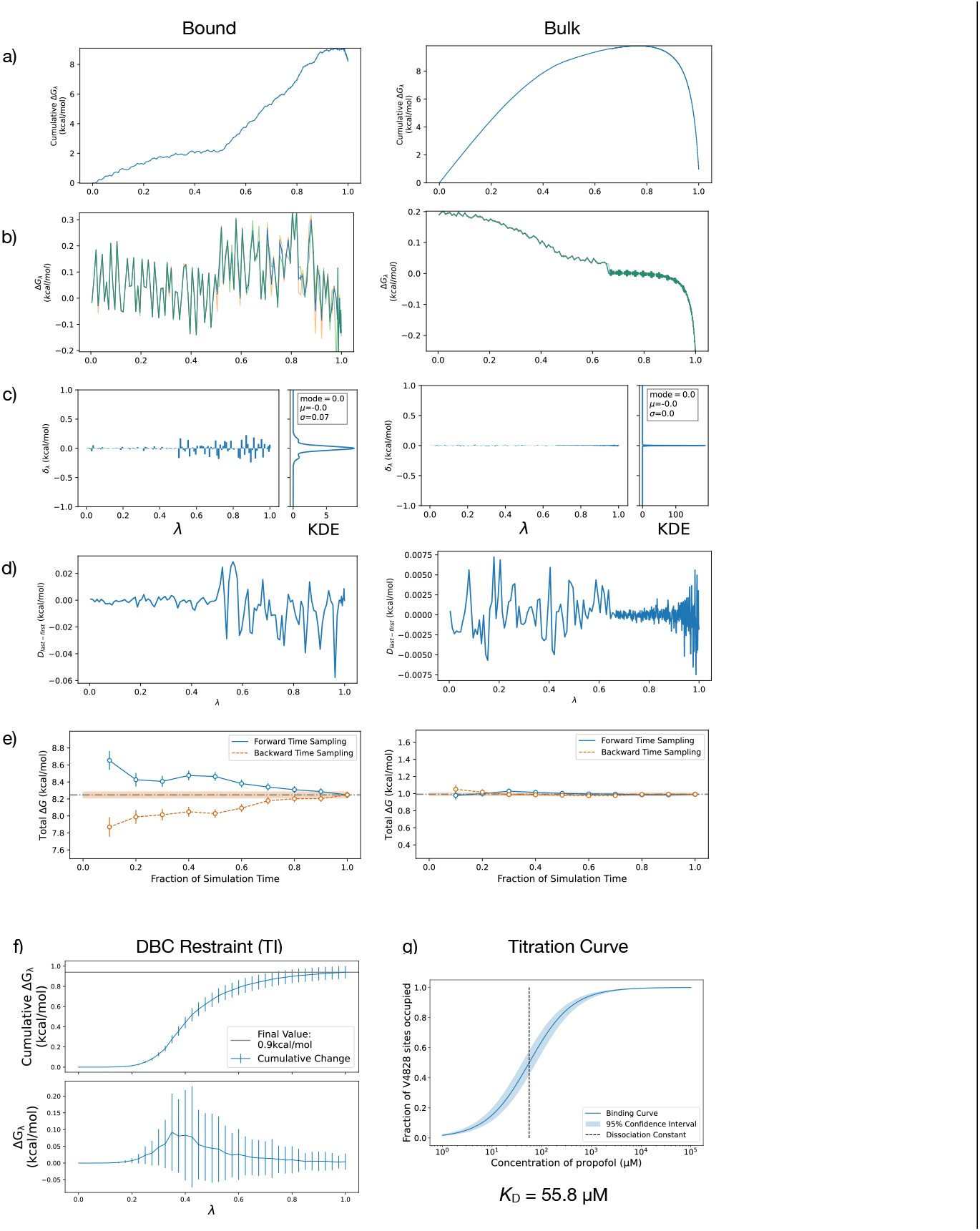
Plots indicating various aspects of convergence of free energy MD simulations. For panels a through e, protein bound state decoupling on left and bulk aqueous phase decoupling on right. a) Cumulative sum of ΔG for each window. b) Per-window ΔG, not summed. c) Discrepancy in ΔG for each window, between forward (λ increasing) and backward (λ decreasing) directions. A kernel density estimation (KDE) of the probability distribution of these values is shown on the right portion of this subpanel. Smaller values imply better convergence. d) Discrepancy in ΔG between first and last half of samples for each window. Smaller values imply better convergence. e) Convergence plot, depicting what fraction of simulation time (x-axis) is necessary to achieve a particular magnitude of discrepancy between forward and backward sampling (y-axis). f) Cumulative and individual ΔG for the thermodynamic integration (TI) calculation to calculate the energetic cost of the DBC restraint g) Titration curve, showing fraction of protein sites occupied as a function of propofol concentration

## Notes

### Competing Interest Statement

The authors have declared no competing interest.

